# Genomic and phenotypic analyses suggest moderate fitness differences among Zika virus lineages

**DOI:** 10.1101/2022.02.01.478673

**Authors:** Glenn Oliveira, Chantal B.F. Vogels, Ashley Zolfaghari, Sharada Saraf, Raphaelle Klitting, James Weger-Lucarelli, Carlos O. Ontiveros, Rimjhim Agarwal, Karla P. Leon, Konstantin A. Tsetsarkin, Eva Harris, Gregory D. Ebel, Shirlee Wohl, Nathan D. Grubaugh, Kristian G. Andersen

## Abstract

RNA viruses have short generation times and high mutation rates, allowing them to undergo rapid molecular evolution during epidemics. However, the extent of RNA virus phenotypic evolution within epidemics and the resulting effects on fitness and virulence remain mostly unknown. Here, we screened the 2015-2016 Zika epidemic in the Americas for lineage-specific fitness differences. We engineered a library of recombinant viruses representing twelve major Zika virus lineages and used them to measure replicative fitness within disease-relevant human primary cells and live mosquitoes. We found that two of these lineages conferred significant *in vitro* replicative fitness changes among human primary cells, but we did not find fitness changes in *Aedes aegypti* mosquitoes. Additionally, we found evidence for elevated levels of positive selection among five amino acid sites that define major Zika virus lineages. While our work suggests that Zika virus may have acquired several phenotypic changes during a short time scale, these changes were relatively moderate and do not appear to have enhanced virulence or transmission during the epidemic.

## Introduction

Zika virus is a mosquito-borne flavivirus that was introduced into the Western Hemisphere in 2014 ^1^, where it infected an estimated 100 million people ^2^. Its genome is translated into a ~3,420-amino acid polyprotein that is processed to yield three structural genes (C, prM, E) and seven non-structural genes (NS1, NS2A, NS2B, NS3, NS4A, NS4B, NS5). Previously thought to cause mild, self-limiting disease, recent discoveries have implicated Zika virus in long-term persistence and pathology in adults ^3–5^. One study demonstrated that seven percent of newborns exposed to Zika virus during pregnancies lead to microcephaly and an additional six percent suffer neurological or ocular defects ^6^. Because these pathologies had not been documented prior to 2015, it has been hypothesized that the virus acquired mutations that enhanced virulence ^7–9^. Additionally, it is possible that the virus adapted to mosquitoes or humans, which may have facilitated its explosive spread across the Americas ^7,8^. These phenomena raise the question of whether Zika virus evolved phenotypically to facilitate spread and/or virulence throughout the Americas.

Several studies have identified putative changes in transmissibility or virulence by studying nonsynonymous mutations that define major Zika virus phylogenetic branches (clades or lineages). For example, Liu et al. found that the NS1-A188V amino acid substitution resulted in higher infectivity of *Aedes aegypti* mosquitoes ^10^, and Shan et al. identified an amino acid substitution, E-V473M, that appears to increase Zika virus virulence and fitness in mice ^11^. Moreover, Yuan et al. showed the prM-S17N amino acid substitution enhanced neurovirulence in mice and replicative fitness in human neural progenitor cells ^12^. They speculated that this mutation may have resulted in an increase in microcephaly incidence ^12^. However, more recent studies have not corroborated this increase in fitness and neurovirulence ^11,13^.

Phenotypic evolution during other RNA virus epidemics further supports the hypothesis that Zika virus may have evolved functional changes during the 2015-2016 epidemic. For example, a chikungunya virus mutation in the envelope gene, E1-A226V, was found to enhance vector competence in *Aedes albopictus* mosquitoes ^14^. Researchers also associated the Ebola virus A82V glycoprotein amino acid substitution with enhanced infection of human cells ^15,16^, and they hypothesized that A82V may have led to increased transmissibility of the virus during the 2015-2016 epidemic in West Africa ^15,16^. More recently, it has been shown that variants of SARS-CoV-2, such as Alpha and Delta, confer higher transmissibility and rapidly displaced other variants across the world ^17–19^. Despite these examples of phenotypic evolution, there has yet to be a genome-wide, systematic screen for phenotypic virus evolution during an epidemic.

To investigate whether Zika virus evolved altered virulence or transmissibility, we screened for fitness differences among lineages that emerged during the 2015-2016 Zika epidemic. We found that the nonsynonymous mutations that define two lineages conferred increased replicative fitness in human primary cells, but we did not find clear differences in lineage-specific fitness in Ae. aegypti mosquitoes. We also found evidence for elevated positive selection among major lineage-defining amino acid sites. However, none of our Zika virus lineages with enhanced replicative fitness displaced ancestral lineages during the epidemic, as has previously occurred during epidemics caused by other viral pathogens ^15,20^. Taken together, our findings suggest that while Zika virus likely acquired phenotypic changes during the 2015-2016 epidemic as it evolved in response to novel environments in the Americas, it is unlikely these changes had a significant impact on the course of the epidemic.

## Results

### Thirteen major Zika clades defined by lineage-specific mutations

To screen for phenotypic evolution among Zika virus lineages during the 2015-2016 epidemic, we identified candidate lineages that may have arose due to adaptive evolution. The candidate lineages were identified using two main criteria: First, because nonsynonymous mutations are more likely to confer phenotypic effects ^21^, we required candidate clades to be defined by at least one nonsynonymous mutation. Second, because phenotypic changes that are adaptive are more likely to proliferate and seed major lineages, we focused on the largest branches within the Zika virus phylogeny. Using these criteria, we selected thirteen major clades that were defined by at least one nonsynonymous mutation.

Specifically, we downloaded 517 unique Zika virus genomes with >90% coverage from Genbank ^22^ that were collected from 2013 to 2019 across Asia and the Americas. Next, we used these sequences to build a time-resolved maximum likelihood phylogenetic tree ^23–25^ and we applied our criteria to identify candidate lineages. We identified lineages that were defined by at least one amino acid site and that had a Shannon entropy ^26^ of greater than 0.2, which we used as a measure of diversity. Our screen resulted in thirteen clades, each defined by one to four nonsynonymous mutations for a total of 17 unique mutations (Fig. 1a; Extended Data Table 1). We named the clades derived during the 2015-2016 epidemic in the Americas clades A-J and the clades that preceded American lineages as pre-American (PA) 1-3 (Fig. 1a).

**Fig. 1.**
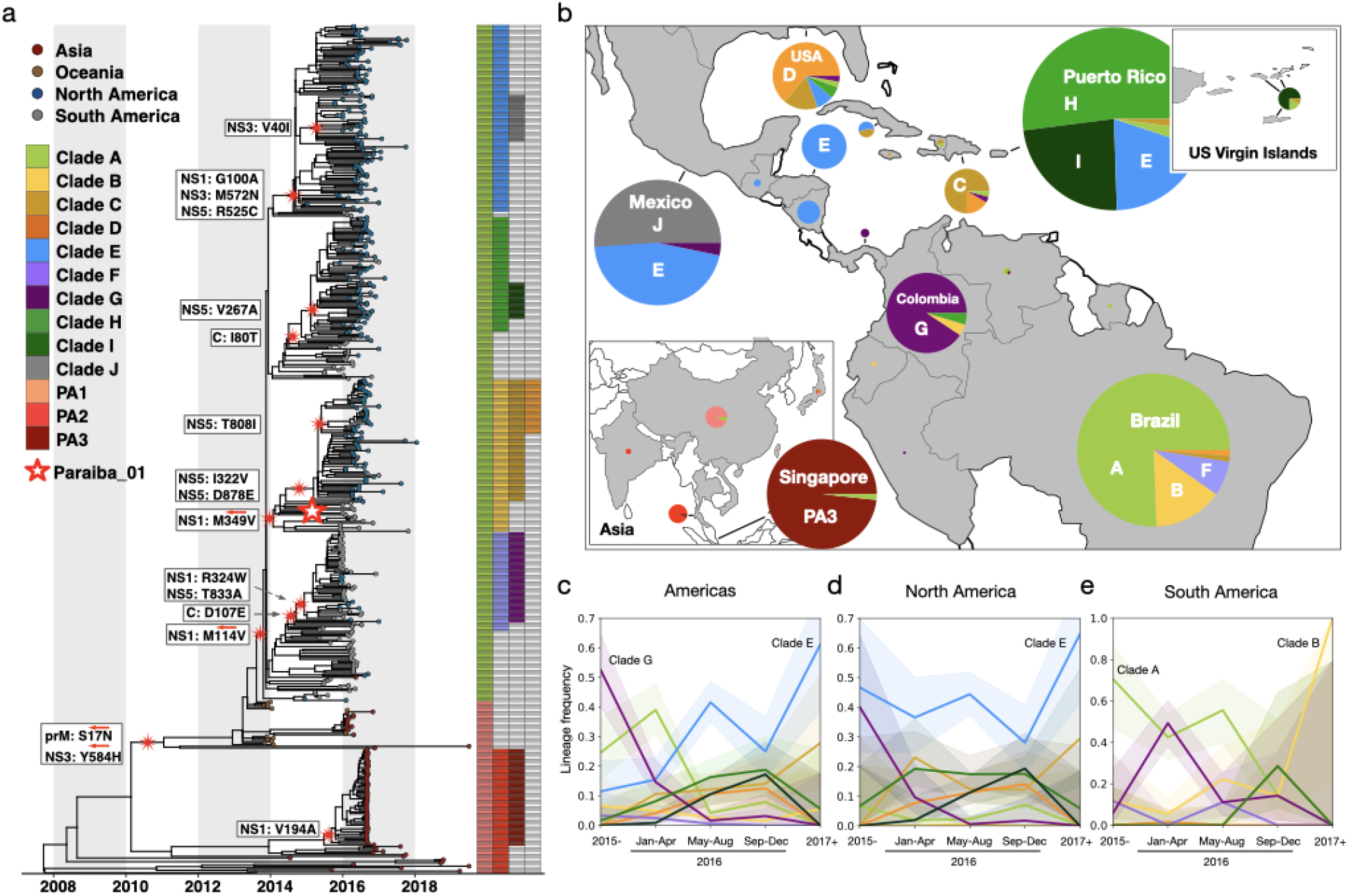
13 major Zika virus lineages defined by nonsynonymous mutations. **(a)**Zika phylogeny partitioned into 13 lineages defined by nonsynonymous mutations. Large red and white star: location of the initial infectious clone into which specific mutations were introduced. White boxes: nonsynonymous mutations introduced into the initial infectious clone to model each lineage. Red arrows: lineage-defining nonsynonymous mutations that were reverted to their ancestral states. **(b)**Proportions of clades circulating in countries across the Americas and Asia. Pie sizes represent the number of sequences. Temporal frequencies of sequenced ZIKV isolates across the Americas **(c)**, North America **(d)**, South America **(e)**with 95% confidence intervals. Timepoint 2015-represents isolates collected during or before 2015. Timepoint 2017+ represents isolates collected during or after 2017.

Instances of adaptive virus evolution are often followed by more fit viral lineages proliferating and displacing ancestral lineages ^15,18,20^. To investigate potential signs of preferential expansion of Zika virus lineages during the 2015-2016 epidemic, we analyzed the temporal and spatial distribution of Zika virus lineages across the Americas, as well as in North and South America individually (Fig. 1b-e). We found the overall frequency of clade E rose 50 and 20 percentage points in the Americas (Fig. 1c) and North America (Fig. 1d), respectively. However, we observed that most lineages did not exceed a total frequency of 70% at any point during the epidemic (Fig. 1c-e). The exception to this is Clade B, which rose to 100% frequency in South America (Fig. 1e); however, this result is based on a single sample sequenced from 2017 or later, suggesting the observed fixation of Clade B is almost certainly due to sampling bias. These findings suggest that while clade E appears to have increased in frequency during the 2015-2016 epidemic, none of these lineages were dominant enough to reach fixation.

### *In vitro* replicative fitness assays in human cells reveal phenotypic differences among Zika virus lineages

To approximate phenotypic evolution during the 2015-2016 epidemic, we investigated the fitness effects of the mutations that define our candidate lineages. We generated Zika virus growth curves by infecting continuous and human primary cells with a library of recombinant Zika viruses that represent 12 Zika virus lineages, and we identified two lineages (clades B and E) that consistently enhanced *in vitro* replicative fitness in human primary cells.

We first used site-directed mutagenesis to introduce the 17 clade-defining mutations into infectious clones that represent our 13 Zika virus lineages (Fig. 1a). We made these mutations on the background of an infectious cDNA clone ^27^ developed from the Paraiba_01 Brazilian Zika virus isolate ^28^, which has been shown to produce viral stocks without introducing additional mutations ^27^. To mitigate potential effects of culture-specific mutations, we transfected each clone into five independent cultures. We successfully rescued infectious clones for 12 of 13 clades (clade J failed to produce infectious virus), leading to a library of 60 viral stocks.

We then used our 60 viral stocks to infect continuous cell lines and human primary cells previously shown to be susceptible to infection ^29–34^. We infected primary human dermal fibroblasts (HDFs), to approximate Zika virus replication at the site of mosquito transmission ^29^, human villous mesenchymal fibroblasts (HVMFs) to recapitulate the ability for Zika virus to disseminate into placental tissue, and human neural progenitor cells (NPCs) to approximate the ability for Zika virus to damage fetal neural tissue. We also infected retinal pigment epithelial (RPE) cells to investigate conjunctivitis ^5,30–32^. To determine whether any phenotype changes were consistent among certain cell types, we also infected continuous cell lines that were similar to our human primary cells (Extended Data Table 2). For all cell types, we generated viral growth curves using plaque assays to measure viral titers across all timepoints (Fig. 2a-h).

**Fig. 2.**
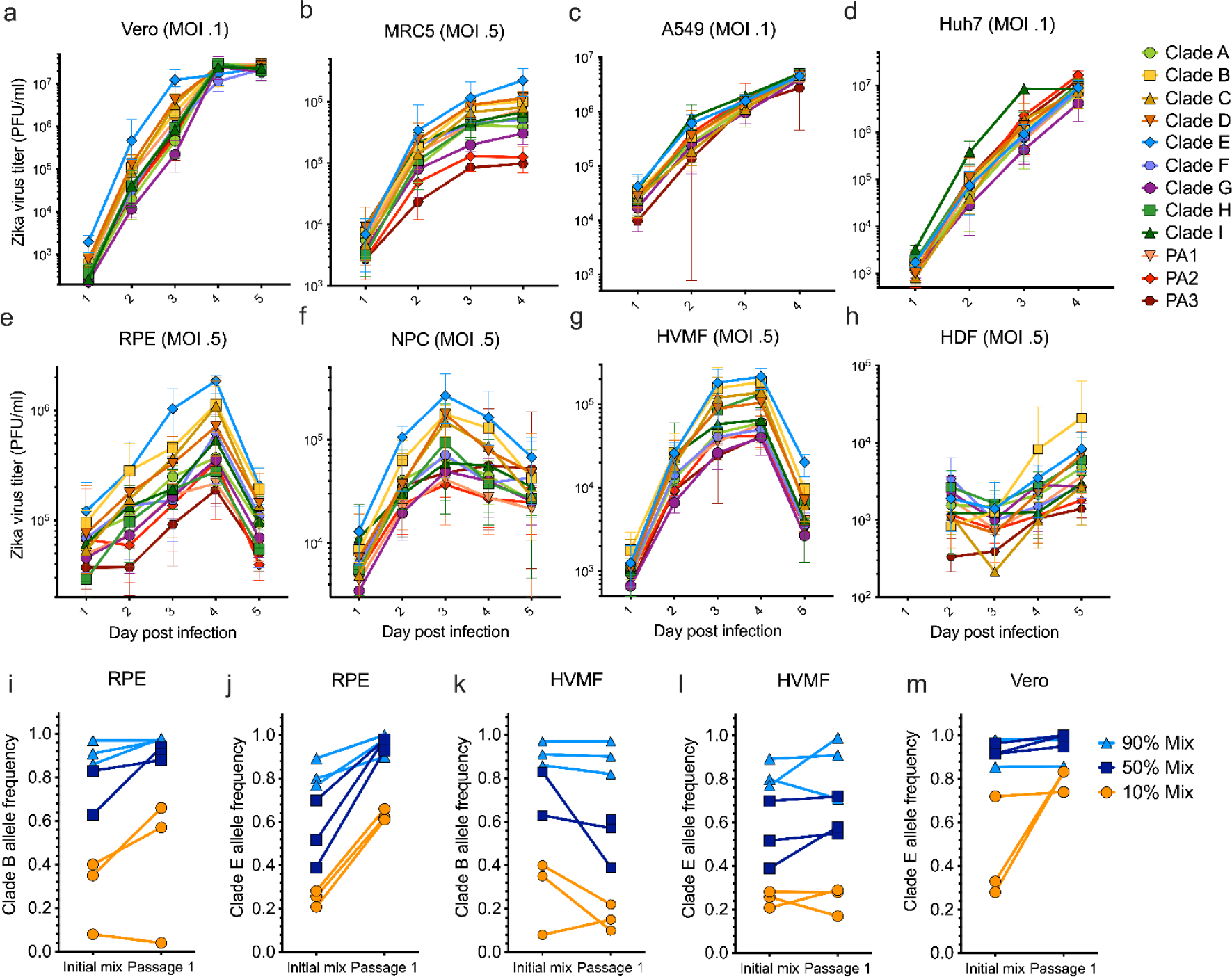
Fitness assays of clade-defining infectious clones in continuous and human primary cells. Replicative fitness of 12 clade-defining recombinant Zika viruses in (**a**) Vero cells, (**b-d**) human continuous cell lines, and (**e-h**) human primary cells. Growth curve points represent the average of the 5 biological replicates and error bars represent their range. Clade B and clade E viruses competed against clade A in (**i, j**) retinal pigment epithelial cells, (**k-l**) human villous mesenchymal cells, and (**m**) the clade E virus competed with clade G in Vero cells. All competitive fitness assays were conducted in triplicate at roughly 90%, 50%, and 10% clade-defining initial frequencies. Vero, African green monkey kidney cells; MRC-5, human lung fibroblasts; A549, human adenocarcinoma alveolar basal epithelial cells; Huh7, human hepatocyte-derived carcinoma cells; RPE, primary human retinal pigment epithelial cells; NPC, human neural progenitor cells; HVMF, primary human villous mesenchymal fibroblasts; HDF, human dermal fibroblasts.

We found that the recombinant virus that represents clade E had the highest and second highest growth over time among all four human primary cells, Vero cells, and MRC-5 cells, and that this clone displayed a significantly higher titer in at least one time point in all of these cell types tested except for HDFs (Fig. 2a-h). Clade E is defined by three amino acid substitutions (NS1-G100A, NS3-M572N, and NS5-R525C; Extended Data Table 1) and spans most of Central America (Fig. 1b). We also found that the clade B recombinant virus had the second highest titers at most timepoints among RPE, NPC and HVMF primary cells (Fig. 2e-2g) and the highest growth curve among HDF primary cells (Fig. 2h). We found the growth curves for the infectious clones that represent clades C and D were similar to clade B within RPE, NPC, and HVMF primary cells (Fig. 2e-2g). Clade B arose in Brazil early in the epidemic and is defined by a single amino acid substitution, NS1-M349V. Clades C and D both descended from clade B (Fig. 1a and 1b) and have other substitutions (NS5-D878E, NS5-T808I) in addition to NS1-M349V (Fig. 1a). We found the replicative fitness curves for clades B, C, and D were clustered in primary cells (Fig. 1a; Extended Data Fig. 1), which suggests that their replicative fitness increases are caused by their shared substitution, NS1-M349V.

We validated the increased replicative fitness of the infectious clones representing clades E and B by performing competitive fitness assays against clade A, the ancestral lineage in the Americas. Each competition was conducted in triplicate at three percentages, 90%, 50%, and 10% for a total of nine competitions per condition. Next, we used deep sequencing to assess lineage frequencies five days post-infection. We infected HVMF and RPE primary cells with these mixed virus populations. Among RPEs, we found that clades E and B increased in frequency in 9/9 and 6/8 mixtures, respectively. Within HVMFs, clade E rose in frequency among 6/9 mixtures but clade B fell in frequency among six of eight mixtures (Fig. 2i-l). To further investigate the high replicative fitness of clade E, we competed it with the lowest performing clade G in Vero cells. Similar to our results with clade A, we found that clade E outcompeted G in 8/9 mixtures (Fig. 2m). These results show that the nonsynonymous mutations that define clades E (NS1-G100A, NS3-M572N, and NS5-R525C), and B (NS1-M349V) confer higher *in vitro* replicative fitness in human primary cells.

### *In vitro* competitive fitness assays using viral isolates confirm replicative fitness differences of recombinant Zika viruses

Above, we used genetically engineered clones to identify lineages with enhanced replicative fitness in human primary cells. To further demonstrate that the clade-defining nonsynonymous mutations specific to clade E and B have high *in vitro* replicative fitness, we investigated if patient-derived Zika virus isolates with these lineage-defining mutations outcompeted isolates with low replicative fitness. Corroborating the fitness results from the engineered infectious clones, we found that virus isolates belonging to clades E and B had enhanced fitness in human primary cells.

We obtained Zika virus isolates from Nicaragua (Nica-6547) and Honduras (R103451) that have shared lineage-defining mutations with clade E, which had the highest replicative fitness in human primary cells (Fig. 2a-h). We also obtained an isolate from Colombia (FLR) and four isolates from Panama (PA259359, PA259634, PAN259249, and PAN259364), that contain the lineage-defining mutations specific to clade G, our lowest-performing clade (Fig. 2a-h). Finally, we included an isolate from Brazil (Paraiba_01) to represent clade B and an isolate from Puerto Rico (PRVABC59) to represent clade I, which we found had a moderate fitness relative to other clones (Fig. 2a-h). We used these isolates to infect HVMF and RPE cells in triplicate to generate *in vitro* replicative fitness curves. In HVMF cells, we found the isolates from Brazil (clade B), Honduras (clade E), and Puerto Rico (clade I) had the highest fitness (Fig. 3a). However, in RPE cells, we found the isolates that represent putative high-fitness clades were clustered with isolates representing clade G and did not demonstrate enhanced fitness (Fig. 3b).

**Fig. 3.**
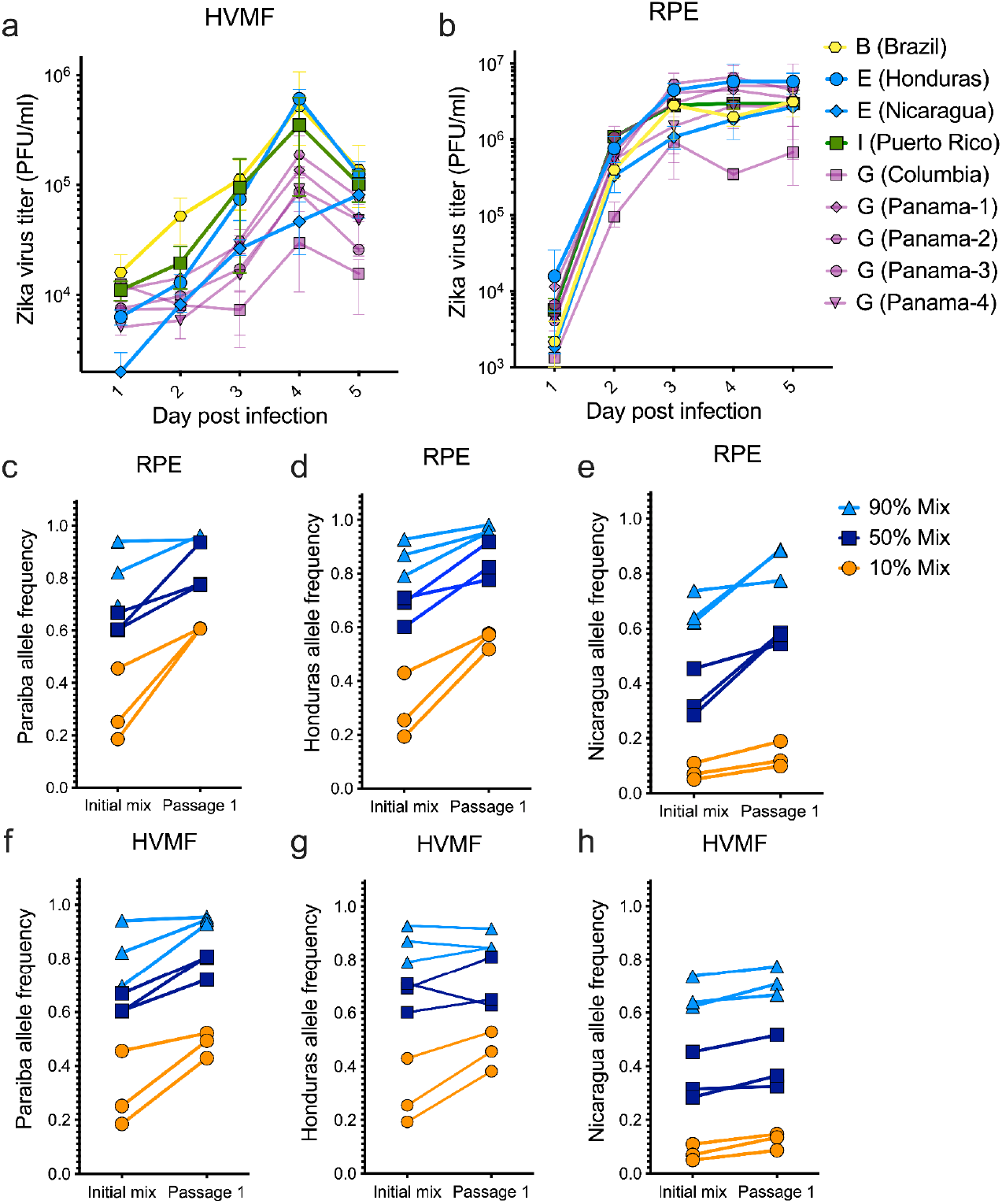
Replicative and competitive fitness results of Zika virus isolates in human primary cells: Infections of (**a**) HVMF and (**b**) RPE cells with nine Zika virus isolates representing clades B, E, G, and I. Growth curve points represent the average of the 5 biological replicates and error bars represent their range. Zika virus isolates from clades with high fitness (clades E and B) competed against a low fitness clade (clade G) using (**c-d**) RPE cells and (**f-h**) HVMF cells. Competitive fitness assays conducted at 90%, 50%, and 10% mixtures and each in triplicate, with three separate isolates representing clade G. RPE, human retinal pigment epithelial cells; HVMF, human villous mesenchymal fibroblasts.

To further investigate fitness effects of our Zika virus isolates, we conducted competitive fitness assays in HVMFs and RPE cells. We competed our clade B isolate (Paraiba_01), and two clade E isolates (Nica-6547 and R103451) against three clade G isolates G (PA259359, PA259634, and PAN259249). Each competition was conducted at three percentages, 90%, 50%, and 10% of each lineage, which led to a total of 27 competitions for each cell type. Finally, we used deep sequencing to assess lineage frequencies five days post infection.

We found that in RPE cells, the Paraiba (clade B), Nicaragua (clade E), and Honduras (clade E) isolates outcompeted clade G isolates (Fig. 3c-e). On average, the increase in frequency rose 21% for Paraiba (clade B), 18% for Honduras (clade E), and 15% for Nicaragua (clade E). In HVMF cells, we found that 24/27 of the competitions resulted in Paraiba (clade B), Nicaragua (clade E), and Honduras (clade E) isolates increasing in frequency (Fig. 3f-h). On average, we found the increase in allele frequencies rose 10% for Paraiba (clade B), 7% for Honduras (clade E), and 5% for Nicaragua (clade E).

We found the HVMF replicative fitness and competitive fitness data suggest enhanced fitness for clades B and E isolates. While we found the RPE competitive fitness data also support enhanced fitness for clades B and E, the isolates that represent these clades did not have elevated replicative fitness curves compared to clade G. Collectively, these findings using Zika virus isolates corroborate the results from our infectious clones demonstrating that clade B and E have elevated replicative fitness in primary human cells.

### Ae. aegypti transmission rates are similar across Zika virus lineages

Zika virus is maintained in a natural transmission cycle between human hosts and mosquito vectors ^35^. To investigate replicative fitness and transmission by mosquitoes, we infected mosquito cells and live mosquitoes with our library of recombinant viruses representing the 12 Zika virus lineages. To inform the selection of mosquito species used for *in vivo* infections, we selected three mosquito cell lines, *Ae. aegypti* Aag2 cells, *Ae. albopictus* U4.4 cells, and *Culex quinquefasciatus* Hsu cells, to represent mosquito species that have been implicated as vectors for Zika virus ^36–38^. We found that both Aag2 and U4.4 sustained Zika virus infection for all clones tested (Fig. 4a and 4b), whereas Zika virus titers rapidly declined in Hsu cells (Fig. 4c), supporting other reports that *Cx. quinquefasciatus* are not competent vectors for Zika virus ^39,40^. Given that *Ae. aegypti* is considered the main vector for Zika virus, and that we observed the highest variation in replicative fitness between clades in these mosquitoes (Fig. 4a), we chose a colony of *Ae. aegypti* from Poza Rica, Mexico ^41^ for *in vivo* fitness evaluations.

**Fig. 4.**
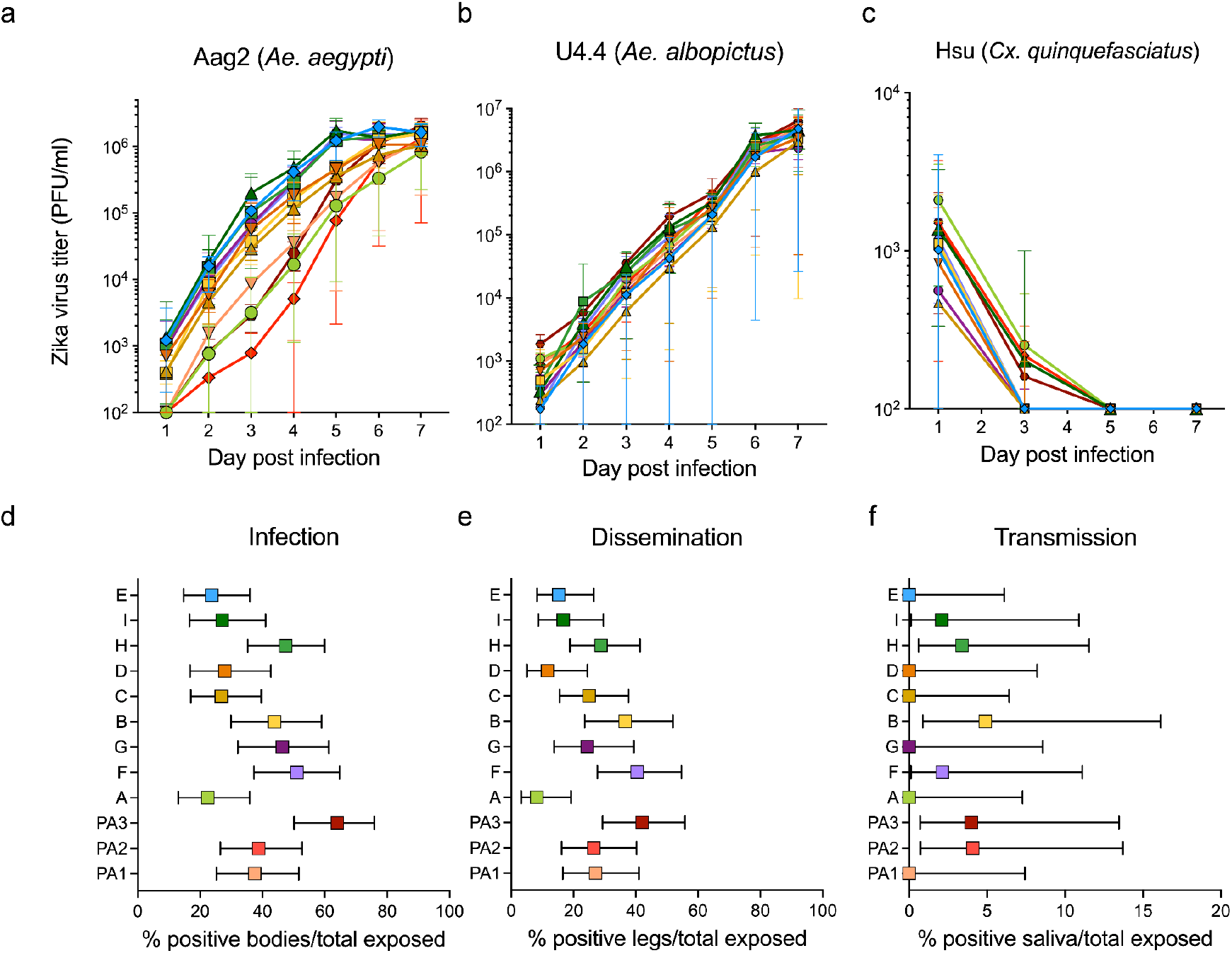
Evaluation of Zika virus fitness in three mosquito cell lines and in *Aedes aegypti* mosquitoes. Replicative fitness of 12 Zika virus clades among infected (**a**) Aag2 (*Aedes aegypti*), (**b**) U4.4 (*Aedes albopictus*), and (**c**) Hsu (*Culex quinquefasciatus*) cells for 3-5 replicates at a multiplicity of infection of 0.1. Squares represent mean titers with error bars depicting the range. *In vivo* fitness of the 12 Zika virus clades by feeding *Aedes aegypti* mosquitoes (Poza Rica, Mexico) with infectious blood meals. (**d**) Infection (percent females with Zika virus-infected body out of the total number exposed females), (**e**) dissemination (percent females with Zika-virus infected legs and wings out of the total number of exposed females), and (**f**) transmission rates (percent females with Zika-virus infected saliva out of the total number of exposed females). Dots represent the rates expressed as percentages and error bars depict the 95% confidence intervals.

To evaluate vector competence of *Ae. aegypti* mosquitoes for each of the 12 clades, we fed groups of female mosquitoes with infectious blood meals containing one of our 12 Zika virus recombinant clones. After 14 days of incubation at 26°C, we dissected mosquitoes and determined the percentage of Zika virus-positive bodies (infection), legs/wings (dissemination), and saliva (transmission) out of the total number of exposed females, via cell culture infectivity assays. We ran pairwise comparisons of infection rates between all clades, and we found a significantly higher infection rate for clade PA3 (Fisher’s exact test with FDR correction, *P* = 0.001) as compared to clade A (Fig. 4d). Furthermore, when running pairwise comparisons of dissemination rates between all clades, we found significantly higher dissemination for clades PA3, B, and F (Fisher’s exact test with FDR correction, *P* = 0.008, 0.03, and 0.008, respectively), as compared to clade A (Fig. 4e). However, we found the subclades nested within clade B (C and D), and clade F (G) do not have elevated dissemination rates (Fig. 4e), which suggests these dissemination differences may not be due to the lineage-defining mutations in question. Lastly, we did not find any significant differences in transmission between the clades (Fisher’s exact test, *P* = 0.28; Fig. 4f). We were not able to detect Zika virus within the saliva of many of our mosquitoes, which likely led to loss of power in our analyses. This may be explained by the relatively low input titer in the blood meal, which was limited by the clade with the lowest titer, to ensure equal titers across all clades.

To further evaluate vector competence, we determined replicative fitness by performing plaque assays on bodies of mosquitoes with and without disseminated infections (*i.e.,* presence/absence of Zika virus in legs and wings). First, we verified Zika virus titers for engorged female mosquitoes that were immediately frozen after feeding. Overall, we found that Zika virus titers were mostly less than 1,000 PFU/mL for all clades, except clade B, which had significantly higher titers than clades A, D, E, G, and I (Dunn’s test with FDR correction, *P* < 0.03; Extended Data Fig. 3a). After 14 days of incubation, we did not detect any significant differences in Zika virus titers for mosquitoes with a disseminated infection when comparing all 12 lineages with each other (Dunn’s test with FDR correction, *P* > 0.05 for all pairwise comparisons; Extended Data Fig. 3b). Thus, although we found that clades PA3, B, and F initially had higher dissemination as compared to clade A, we did not find any differences in replicative fitness associated with the tested lineage-defining nonsynonymous mutations.

Because we found low transmission rates after feeding *Ae. aegypti* females with an infectious blood meal, we wanted to further investigate potential changes in replicative fitness inside mosquitoes. We intrathoracically injected groups of *Ae. aegypti* females with each of our 12 Zika virus clones and determined the percentage of positive saliva (transmission) after seven days incubation. We verified that all injected female mosquitoes, for each clade (Extended Data Fig. 4a), were positive for Zika virus. We compared transmission rates between mosquitos infected with different clones and again found no significant differences between any of the clades (Fisher’s exact test with FDR correction, *P* > 0.05 for all pairwise comparisons; Extended Data Fig. 4b). Taken together, our experiments do not clearly identify lineage-specific fitness differences in live *Ae. aegypti* mosquitoes.

### Clade-specific amino acid sites are under heightened positive selection

Selection pressures among the protein-coding portion of a genome can be analyzed by comparing the rate of nonsynonymous mutations with the rate of synonymous mutations (dN/dS) ^42^. To further evaluate phenotypic evolution during the 2015-2016 epidemic, we calculated dN/dS values ^43^ across all amino acid sites in the Zika virus genome and found 25 amino acid sites that show evidence of positive selection.

Specifically, we applied renaissance counting ^43^ selection analysis to our 517 sequences to calculate site-specific dN/dS values across the protein-coding region of the Zika virus genome ^43^. Among the ~3,420 amino acid sites in the Zika virus genome, we identified 25 sites that had 90% of their posterior dN/dS distributions greater than one (Fig. 5a and 5b). Five of 17 lineage-defining mutations were among the 25 putative sites under positive selection (clades B; NS1-M349V, C; NS5-I322V, G; NS5-T833A, H; C-I80T, and PA2; NS3-Y584H), with average posterior dN/dS ratios ranging from 1.2-1.9 (Fig. 5a and 5b). Throughout the Zika virus genome, 20 amino acid sites that did not define major clades had dN/dS values greater than one, and two of these sites were within two codons from a clade-defining mutation (clades E and PA2). Notably, the NS1-M349V mutation, which defines clade B, was one of two lineage-defining mutations that we found to have a high replicative fitness in human primary cells.

**Fig. 5.**
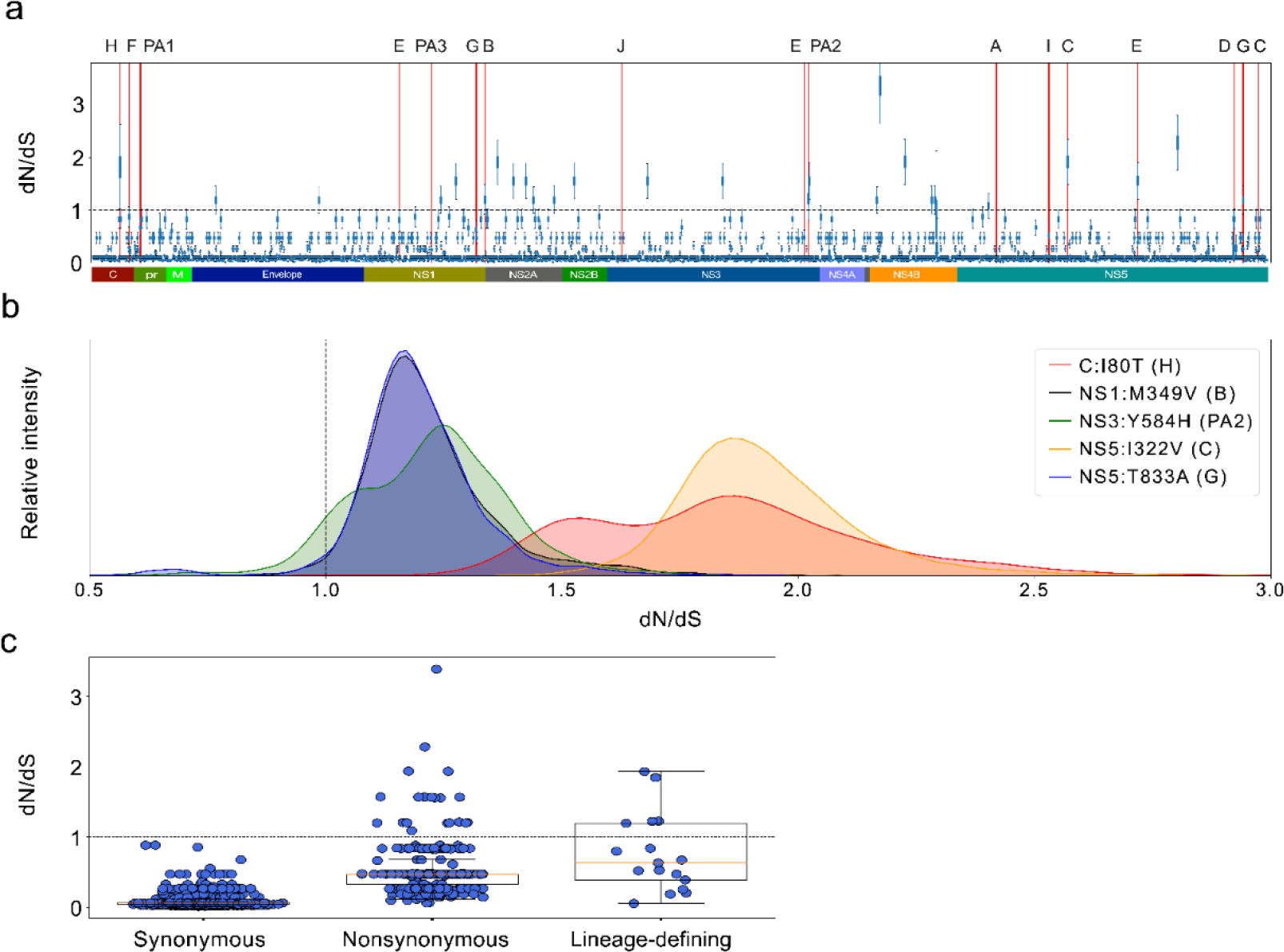
Site-specific dN/dS values across the ZIKV polyprotein. (**a**) Zika virus site-specific dN/dS analysis measured using 517 Zika virus isolates. Box plots: posterior probability distributions of dN/dS values across the Zika virus genome. Red bars: clade-defining loci. (**b**) Posterior probability densities for the five clade-defining amino acid loci that have dN/dS values greater than one. (**c**) The number of synonymous, nonsynonymous, and clade-defining mutations across the Zika virus genome. Clade-specific amino acid sites have significantly higher dN/dS values (p=.0002; Mann-Whitney U test) when compared to the remaining amino acid sites with nonsynonymous mutations.

To investigate whether dN/dS values were significantly higher among lineage-defining mutations as a whole, we compared the dN/dS distributions at lineage-defining amino acids with all nonsynonymous mutations that are not lineage defining. We used ancestral state reconstruction ^24^ to identify all mutations throughout our phylogeny and identified 3,633 synonymous mutations across 1,742 amino acid sites, 825 nonsynonymous mutations across 384 amino acid sites, and 89 nonsynonymous mutations that define our clades across 17 amino acid sites. The average dN/dS values for synonymous only, nonsynonymous, and clade-defining sites were .07, .53, and .76, respectively. Next, we compared the clade-defining amino acid sites with all other nonsynonymous, amino acid-specific dN/dS values, and we found that the clade-defining values were significantly higher (Mann-Whitney U test, p = .04; Fig. 5d and 5e). This shows that lineage-defining amino acid sites tend to have higher dN/dS ratios than non-lineage-defining sites, suggesting that our sites are either under increased positive selection or relaxed purifying selection.

## Discussion

The sudden rise of Zika virus transmission and disease during the 2015-2016 epidemic in the Americas raised the question of whether the virus evolved to become more transmissible or virulent ^7–9^. In this study, we addressed this question by using engineered recombinant viruses representing specific Zika virus lineages to infect human primary cells and live mosquitoes. We found evidence of two lineages with replicative fitness changes in human tissues, but none in Ae. aegypti mosquitoes. Using selection analysis, we found elevated signals for adaptive evolution among five lineages that arose during this time. However, these changes did not result in the displacement of less fit lineages by fitter ones, and the overall phenotypic effects of these lineage-defining mutations warrant further investigation. The ongoing COVID-19 pandemic has demonstrated the ability for a virus to evolve towards higher transmissibility or immune escape during short timescales. This highlights the importance of identifying instances of phenotypic evolution during an outbreak ^44^. Here, we present a systematic approach to identify lineage-specific fitness increases during an epidemic.

We found that the amino acid substitutions specific to clade E (NS1-G100A, NS3-M572N, and NS5-R525C) and clade B (NS1-M349V) confer replicative fitness increases in human primary cells and continuous cells lines. Consistent with our finding that clade E leads to higher Zika virus replicative fitness, Kuo et al. previously demonstrated that the NS1-G100A substitution leads to increased fitness in type-I interferon receptor knockout (A129) mice ^47^. NS1-G100A is one of the three amino acid substitutions that define clade E, and it is plausible this substitution confers some, or all, of the fitness advantage in the human primary cells we tested. While further work will be required to decipher which of these three mutations leads to higher replicative fitness in human cells, our and Kuo et al.’s analysis are suggestive that clade E confers a fitness advantage in mammalian hosts.

The lack of microcephaly observed before Zika virus spread to Oceania and the Americas raised the possibility that Zika may have evolved towards increased virulence. Yuan et al. ^12^ found the prM-S17N substitution increased fitness in neural progenitor cells and conferred shorter survival when injected intracranially into one-day-old mice. They hypothesized that this substitution enhanced microcephaly incidence or severity. However, consistent with Jaeger et al.’s and Shan et al.’s analysis, we did not find evidence to support this theory. Specifically, Shan et al. demonstrated that prM-17N and prM-17S have nearly identical survival curves when intracranially infecting one-day-old mice, suggesting the prM-17N does not confer enhanced disease. Additionally, Jaeger et al. did not identify fitness or neurovirulence differences when infecting these prM-17 mutants in one-day-old mice. In our analysis, the prM-S17N substitution occurs between lineages PA2 and clade A, and in contrast to Yuan et al. ^12^, we did not identify any fitness changes between these lineages in neural progenitor cells. However, our experiments also included the NS3-Y584H mutation, which co-defines the PA2 lineage, so we were not able to measure the effect of the prM-S17N substitution independently.

Because we assessed replicative fitness differences from *in vitro* infections of human primary cells and laboratory mosquitoes, it is difficult for us to determine if any lineage enhanced Zika virus fitness in nature. In our experiments, we used NPCs, HVMFs, and RPEs to model disease and HDFs to model transmission. While we selected these cell types because they have been shown to be susceptible to Zika virus ^5,29–31^, they may not be the best models to approximate these phenotypes. For example, human monocytes likely play a role in Zika virus transmission ^45,46^ but we were unable to infect these cells at high enough levels to generate growth curves. Nonetheless, we found a strong concordance of replicative fitness differences among the human primary cells included in our analyses (Fig. 2a-h and Extended Data Fig. 1), suggesting that the fitness differences we identify likely confer similar effects across a broad range of human tissues.

To bolster our replicative fitness results in specific model systems, we investigated the selective pressures across the Zika virus genome. We found evidence that five lineage-defining amino acid sites may be under positive selection and that sites that define major lineages have significantly elevated dN/dS values. Notably, we found evidence for positive selection at the NS1-M349V amino acid site that defines clade B, which we also found to have high replicative fitness in human primary and continuous cells. While our selection analysis is valuable in the context of our replicative fitness assays, these data should be interpreted with caution. First, dN/dS analysis is prone to underestimate positive selection during viral epidemics due to little time for purifying selection purge to deleterious mutations ^42^. Additionally, this analysis assumes that synonymous mutations are mostly neutral, which is not always the case for RNA viruses ^48^. Therefore, we used our selection analysis as a screen to identify candidate sites for positive selection and to support our replicative fitness findings.

Our selection analysis may also enable us to investigate functional differences across a full transmission cycle. Because we evaluated Zika virus fitness in humans and mosquitoes separately, we could not evaluate how replicative fitness changes in either organism would affect overall fitness in a complete, natural transmission cycle. Our dN/dS analysis, however, incorporates the full evolutionary history in mosquitoes and humans. Hence, it is possible that a subset of the five lineage-specific mutations with high dN/dS values confer fitness advantages in a natural transmission cycle. Additionally, we note that the two lineages that appear to confer fitness advantages in human primary cells (clades B and E) do not appear to be deleterious in live mosquitoes or the Aag2 mosquito cell line, suggesting that these fitness increases could remain during a full transmission cycle.

While our study identifies multiple lineage-specific phenotypic changes during the course of the Zika epidemic, these changes did not result in the fixation of their respective lineages, and therefore are unlikely to have had major effects on the trajectory of the epidemic. It is possible that the lineages we found to have enhanced fitness in human primary cells may have been tempered by low or moderate fitness in mosquitoes. Moreover, consistent with a rapid decline in Zika cases after the 2015-2016 epidemic ^49^, none of our 13 major lineages were defined by mutations in the envelope gene. The envelope protein is responsible for host-receptor binding, and the lack of lineages defined by amino acid substitutions in this gene suggests immune escape variants were not able to proliferate to high frequencies. Therefore, it is unlikely that antibody-specific immune evasion significantly impacted this epidemic.

Here, we provide a framework to screen for fitness differences among lineages during an epidemic. While the fitness changes we identified do not appear to have significant effects on the 2015-2016 Zika epidemic, our analysis suggests phenotypic evolution occurred several times during this outbreak. Monitoring the phenotypic evolution during the course of an outbreak can help control spread and mitigate disease ^44^. We believe this framework can be applied to study phenotypic evolution during future epidemics caused by emerging RNA viruses.

## Methods

### Cell lines and viruses

Vero (ATCC:CCL-81), Huh-7, A549 (ATCC), SH-SY5Y, and MRC5 (ATCC) cell lines were maintained in 1X Dulbecco’s modified eagle medium (Gibco) supplemented with 10% heat-inactivated fetal bovine serum, 1mM sodium pyruvate (Gibco), and 1x penicillin-streptomycin (Gibco), and were grown at 37°C with 5% CO2. RPE, HDF, and HVMF human primary cells and respective media were purchased from Sciencell and the manufacturer’s instructions were followed for culturing. NPC human primary cells and supporting culture ingredients were purchased from STEMCELL technologies and the manufacturer’s instructions were followed.

We used mosquito-derived cell lines from three mosquito species to investigate replicative fitness of the twelve Zika virus mutants. We inoculated twelve Zika virus mutants on *Ae. aegypti*-derived Aag2 cells, *Ae. albopictus*-derived U4.4 cells, and *Cx. quinquefasciatus*-derived Hsu cells (kindly provided by Dr. Doug Brackney, Connecticut Agricultural Experiment Station). We maintained Aag2 cells in Schneider’s Drosophila Medium supplemented with 8% heat-inactivated fetal bovine serum and 1% antibiotic/antimycotic (100 U/mL penicillin, 100 μg/mL streptomycin, and 0.25 μg/mL amphotericin B) at 28°C without CO2. U4.4 cells were maintained in Mitsuhashi and Maramorosch insect medium supplemented with 7% heat-inactivated fetal bovine serum, 1% antibiotic/antimycotic, 1% non-essential amino acids, and 1% L-glutamine at 28°C with 5% CO2. We maintained Hsu cells in 1X minimum essential Earle’s medium supplemented with 10% heat-inactivated fetal bovine serum, 1% antibiotic/antimycotic, and 1% non-essential amino acids at 28°C with 5% CO2. For infectivity and plaque assays, we used Vero E6 cells (African green monkey; kindly provided by Doug Brackney, Connecticut Agricultural Experiment Station), which we maintained in 1X minimum essential Earle’s medium supplemented with 10% heat-inactivated fetal bovine serum, 1% antibiotic/antimycotic, and 1% non-essential amino acids at 37°C with 5% CO2.

We obtained the FLR, R103451, PAN259249, PAN259634, PAN259359, PAN259364 viral isolates through BEI Resources, NIAID, NIH as part of the Human Microbiome Project. Nica-6457 was isolated from a participant of the Pediatric Dengue Cohort Study in Managua, Nicaragua, and was kindly provided by Eva Harris (University of California, Berkeley), and Paraiba_01 isolate was kindly provided by Thomas Rogers (Scripps Research, La Jolla).

### Generating clade-defining infectious clones

We used the phylogeny and diversity data on Nextstrain ^50^ to select major clades that are defined by nonsynonymous mutations, where the amino acid site had a Shannon entropy of greater than 0.2 as of February 2020. We used an *in vitro* assembly mutagenesis method to introduce clade-defining mutations into a Zika virus infectious clone generated from the Brazilian isolate Paraiba_01 kindly provided by Dr. Alexander Pletnev ^27^. Next, we sequenced these plasmids on an Illumina Miseq to confirm the addition of all mutations and omit infectious clones with minor allele frequencies above 20% at unintended sites. We then transfected 3 ug of these infectious clones into Vero cells at ~75% confluence in a T25 flask by using lipofectamine 3000. The transfections were repeated five times for each infectious clone to control for spurious effects arising during transfection and culture. We rescued virus stocks on day three and partitioned into 24 100ul aliquots, where Hepes buffer was added to a final concentration of 1%. We used triplicate plaque assays to measure the titer of each viral stock. The plaque assays were conducted in 96-well plates with Vero cells seeded to a ~80% confluency. The virus stocks were serially diluted from 100 to 100,000-fold. The diluted virus was incubated with Vero cells for 1.5 hours before the addition of a 2% methylcellulose/DMEM mixture. The infected Vero cells were then incubated at 37°C and 5% CO_2_ for five days.

### Mosquitoes

We maintained *Ae. aegypti* mosquitoes (F24-26) originating from Poza Rica, Mexico in bugdorm-1 screen mesh cages at 27°C, 60-70% relative humidity, and 12:12 light-dark cycle. Adult mosquitoes were maintained on 10% sucrose solution and females were blood-fed with defibrinated sheep blood. Eggs were transferred to rearing trays containing 500 mL of water and two drops of Liquifry No. 1. Approximately 250 hatched L1 larvae were transferred to a tray with one L of water and two drops of Liquifry No. 1. Larvae were fed with tetramin baby fish food. Females were transferred to 32 oz soup cartons, screened with fine mesh for Zika virus infections in the BSL2 insectary. One day before infectious blood meals were provided, sucrose solution was replaced with water.

### Mosquito infections

To determine Zika virus fitness *in vivo*, we exposed groups of 7-12 day-old female *Ae. aegypti* mosquitoes to each of the 12 Zika virus mutants in parallel, for a total of three independent biological replicates. Zika virus was mixed with defibrinated sheep blood to a final titer of 7.5E5 PFU/mL and provided through the Hemotek artificial membrane feeding system using a parafilm membrane. We allowed female mosquitoes to feed for approximately one hour, and then immobilized them on ice to select engorged females. We counted engorged females and placed them in a new carton closed with a fine mesh and provided access to cotton wool soaked in 10% sucrose solution. After 14 days of incubation at 26°C, we immobilized female mosquitoes on ice and removed legs and wings. We collected legs and wings in a safe-lock tube with a steel bead and 200 μL of mosquito diluent. Mosquito diluent consisted of 1X phosphate buffered saline with 20% fetal bovine serum, 50 μg/mL penicillin/streptomycin, 50 μg/mL gentamycin, and 2.5 μg/mL amphotericin B. We inserted the proboscis into a 200 μL pipette tip containing five μL of a 1:1 solution of fetal bovine serum and 50% sucrose solution to collect saliva for approximately one hour. After one hour, we transferred saliva to tubes containing 100 μL of mosquito diluent, and we transferred mosquito bodies to a safe-lock tube with a steel bead and 200 μL of mosquito diluent.

In addition to exposing female mosquitoes to the twelve Zika virus mutants via an infectious blood meal, we also injected each of the mutants in the thorax. We used the nanoject two auto-nanoliter injector to inject 69 nL of 1.5E6 PFU/mL in each female mosquito, for three independent biological replicates. We injected mosquitoes and waited for seven days, until saliva was collected as described above. We tested both mosquito bodies and saliva for presence of Zika virus via infectivity assays.

### Replicative Fitness assays

We used our library of 60 viral stocks to infect seven continuous cell lines and four human primary cell types. A multiplicity of infection (MOI) of 0.5 was used to infect all primary lines and MRC5s and an MOI of 0.1 was used to infect the remaining continuous cell lines. After dilution to MOI of 0.1 or 0.5, the inoculum was added to a 24-well plate well seeded with ~100,000 cells and incubated for one hour. Next, the inoculum was removed, and the cells were washed once with assay media. Finally, 0.5 mls of assay medium was added and 50 ul was sampled every 24 hours with replacement of assay media. We used triplicate plaque assays to measure viral titers at each time point.

### Competitive Fitness Assays

For competitive fitness assays, we used PFU titers to mix two viral mutants at three ratios: 90:10, 50:50, and 10:90. We used these inocula to infect ~100,000 Vero cells in triplicate. We incubated the inoculum in the 24-well plate for one hour and washed once with assay medium. After five days, 100ul was set aside for sequencing and another 100ul was used to inoculate a fresh well of Vero cells. This process was repeated for a total of three passages. The banked samples were deep sequenced using an Illumina Miseq, and the allele frequencies for each mutant were measured using iVar ^51,52^. This process was repeated for primary cells.

### Mosquito infectivity assays

We used infectivity assays to determine whether mosquito parts and saliva were infected with Zika virus. We seeded Vero E6 cells in 96-wells the day before inoculations. We blended mosquito body parts in the Bullet blender storm 24 for two minutes at maximum speed (speed 10). We centrifuged all samples (including mosquito saliva) for two minutes at 13,000 rpm. We removed medium from 96-well plates and added 50 μL of sample to each well. After 2-3 hours incubation, we removed the supernatant and added 100 μL of MEM supplemented with 10% fetal bovine serum, 1% antibiotic/antimycotic, 1% non-essential amino acids, 50 μg/mL gentamicin, and 2.5 μg/mL amphotericin B to each well. We incubated the plaque assay plates at 37°C with 5% CO2 and we screened for cytopathic effects after seven days.

### Evolutionary analysis

We downloaded the Zika virus consensus sequences from the ViPR database ^53^, and omitted all sequences that spanned less than 90% of the Zika virus genome. We used MUSCLE ^23^ combined all unique sequences with our previous Zika virus multiple sequence alignment reported in our previous publications that resulted in an alignment of 517 unique Zika virus consensus sequences. We used IQ-TREE ^25^ to generate a maximum likelihood tree (Fig. 1A) and treetime ^24^ to date ancestral nodes. For temporal and spatial profiling of the American lineages, we generated a maximum likelihood tree as described above, but we used 579 unique Zika virus consensus sequences.

To estimate dN/dS values across our Zika virus phylogeny, we used Renaissance counting ^43,54^. Specifically, we used the coding region of our 517 Zika virus sequence alignment to conduct this analysis. We used an HKY nucleotide substitution model ^55^ with all three codon positions partitioned. We used an uncorrelated relaxed log normal clock ^56^ that we determined to be the ideal clock rate for Zika virus phylogeny in previous studies ^1,57^. We ran this analysis in BEAST for 250 million states while sampling every 10,000 states. We discarded the initial 25 million states. To determine which of the 3423-amino acid sites in the Zika virus genome contained synonymous only, and at least one nonsynonymous changes, we used ancestral state reconstruction in TreeTime ^24^.

### Statistical analysis

To screen the mammalian Zika virus growth curves for fitness differences, we applied the Mann-Whitney U test at individual time points, and we used FDR to correct for multiple comparisons ^58,59^. These analyses were conducted using the SciPy and Statsmodels package in Python ^60,61^. To analyze the mosquito experiments, we used Fisher’s exact tests to test for significant changes in infection, dissemination, and transmission, with FDR correction to account for multiple comparisons. To test for differences in virus titers, we used Kruskall-Wallis tests, and Dunn’s post-hoc test with FDR correction for multiple comparisons ^62^. Statistical analyses were done in R version 3.6.1.

To allow comparisons of fitness values across cell types and mosquito models, we averaged the day three and day four (day three for A549 cells) replicative fitness values and converted them to Z-scores. The Z-scores for each model were clustered (agglomerative hierarchical clustering) using the seaborn package in python ^63^ (Supplementary Fig. 4)^3^.

## Acknowledgements

We thank Mark Zeller, Gytis Dudas, Edyth Parker, Karthik Gangavarapu and Nathaniel Wineinger for insightful discussions. Thank you to Refugio Robles-Sikisaka for laboratory support. We thank Alexander Pletnev for generously providing the infectious Zika virus clone. We thank Dr. Angel Balmaseda, Dr. Guillermina Kuan, and the entire study team of the Pediatric Dengue Cohort Study in Managua, Nicaragua, which provided the Nicaraguan isolate Nica-6547, supported by NIH grants P01AI106695 (E.H.), U19AI118610 (E.H.), and R01AI099631 (A.B.). C.B.F.V. is supported by NWO Rubicon (no. 019.181EN.004). K.G.A. is a Pew Biomedical Scholar and is supported by NIH NCATS CTSA UL1TR002550, NIAID contract HHSN272201400048C, NIAID R21AI137690, NIAID U19AI135995, and The Ray Thomas Foundation.

## Data availability

Illumina sequence data are publicly available on the SRA database under BioProject PRJNA788964. Zika virus sequences are publicly available on NCBI’s Genbank. Genbank accession codes, plaque assay, and dN/dS data are available at: https://github.com/andersen-lab/paper_2022_zika-functional-evolution.

## Code availability

Sequence data was processed using iVar software ^52^. Custom scripts are available at https://github.com/andersen-lab/paper_2022_zika-functional-evolution.

## Supplementary data

**Extended Data Table 1.** Lineage-defining nonsynonymous mutations

| Extended Data Table 1 \| Lineage-defining nonsynonymous mutations |  |  |  |  |
| --- | --- | --- | --- | --- |
| Clade | Gene | Gene location | Ancestral amino acid | Mutant amino acid |
| Clade A | NS1 | 349* | M | V |
| Clade B | No changes needed, Paraiba_01 is this genotype |  |  |  |
| Clade C | NS5 | 322 | I | V |
|  | NS5 | 878 | D | E |
| Clade D | NS5 | 322 | I | V |
|  | NS5 | 808 | T | I |
|  | NS5 | 878 | D | E |
| Clade E | NS1 | 349* | M | V |
|  | NS1 | 100 | G | A |
|  | NS3 | 572 | M | L |
|  | NS5 | 525 | R | C |
| Clade F | NS1 | 349* | M | V |
|  | Capsid | 107 | D | E |
| Clade G | NS1 | 349* | M | V |
|  | Capsid | 107 | D | E |
|  | NS1 | 324 | R | W |
|  | NS5 | 833 | T | A |
| Clade H | C | 80 | I | T |
|  | NS1 | 349* | M | V |
| Clade I | Capsid | 80 | I | T |
|  | NS1 | 349* | M | V |
|  | NS5 | 267 | V | A |
| Clade J | NS1 | 349* | M | V |
|  | NS1 | 100 | G | A |
|  | NS3 | 572 | M | L |
|  | NS3 | 40 | V | I |
|  | NS5 | 525 | R | C |
| PA1 | NS1 | 349* | M | V |
|  | NS5 | 114* | M | V |
| PA2 | NS1 | 349* | M | V |
|  | prM | 17* | S | N |
|  | NS3 | 584* | Y | H |
|  | NS5 | 114* | M | V |
| PA3 | NS1 | 349* | M | V |
|  | NS5 | 114* | M | V |
|  | prM | 17* | S | N |
|  | NS3 | 584* | Y | H |
|  | NS1 | 194 | V | A |

\*Paraiba\_01 infectious clone was reverted to its ancestral amino acid state.

**Extended Data Table 2.** Experimental Cell types

| Extended Data Table 2 \| Experimental Cell types |  |  |  |  |
| --- | --- | --- | --- | --- |
| Cell type | Organism | Tissue type | Description | Prior Zika virus studies |
| HDF | Human | Fibroblast | Human Dermal fibroblasts | Hamel et al. |
| HVMF | Human | Fibroblast | Human villous mesenchymal fibroblasts | El Costa et al. |
| RPE | Human | Epithelial | Retinal pigment epithelial cells | Manangeeswaran et al. |
| NPC | Human | Neural | Neural progenitor cells | Li et al. |
| Vero | African green monkey | Epithelial | Kidney epithelial cells | Vicenti et al. |
| MRC5 | Human | Fibroblast | Fibroblasts derived from lung tissue | Vicenti et al. |
| A549 | Human | Epithelial | Adenocarcinomic alveolar basal epithelial cells | Vicenti et al. |
| Huh7 | Human | Epithelial | Hepatocyte-derived carcinoma cell line | Vicenti et al. |
| SHSY5Y* | Human | Neural | Neuroblast derived from bone-marrow cell line | Luplertlop et al. |
| Aag2 | *Ae. aegypti* | Mixed | Derived from homogenised embryos | Weger-Lucareli et al., Chouin-Carneiro et al |
| U4.4 | *Ae. albopictus* | Unknown | Derived from neonate larva | Chouin-Carneiro et al |
| Hsu | *Cx. quinquefasciatus* | Unknown | Derived from ovarian tissue | Guo et al., Weger-Lucareli et al. |

\*SHSY5Y infections yielded detectable virus during only 1 timepoint and are not included in figure 2.

**Extended Data Fig. 1.**
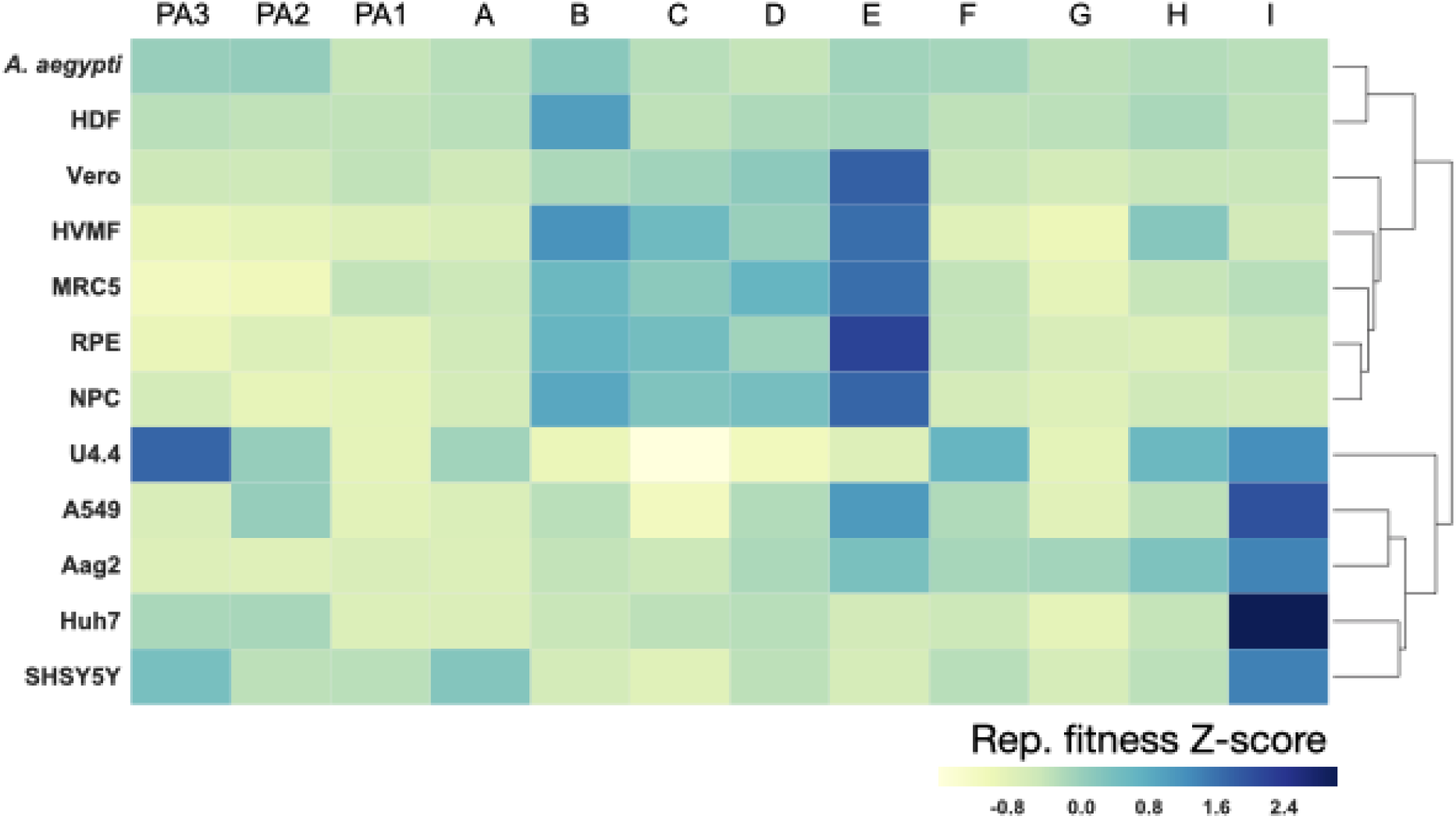
Hierarchical clustering of model-specific fitness values. To compare fitness values across cell and mosquito models, we averaged the day three and day four (just day three for A549) replicative fitness values and converted them to Z-scores. Then the Z-scores for each model were clustered (agglomerative hierarchical clustering). Cell types are explained in (**Extended Data Table 2**).

**Extended Data Fig. 2.**
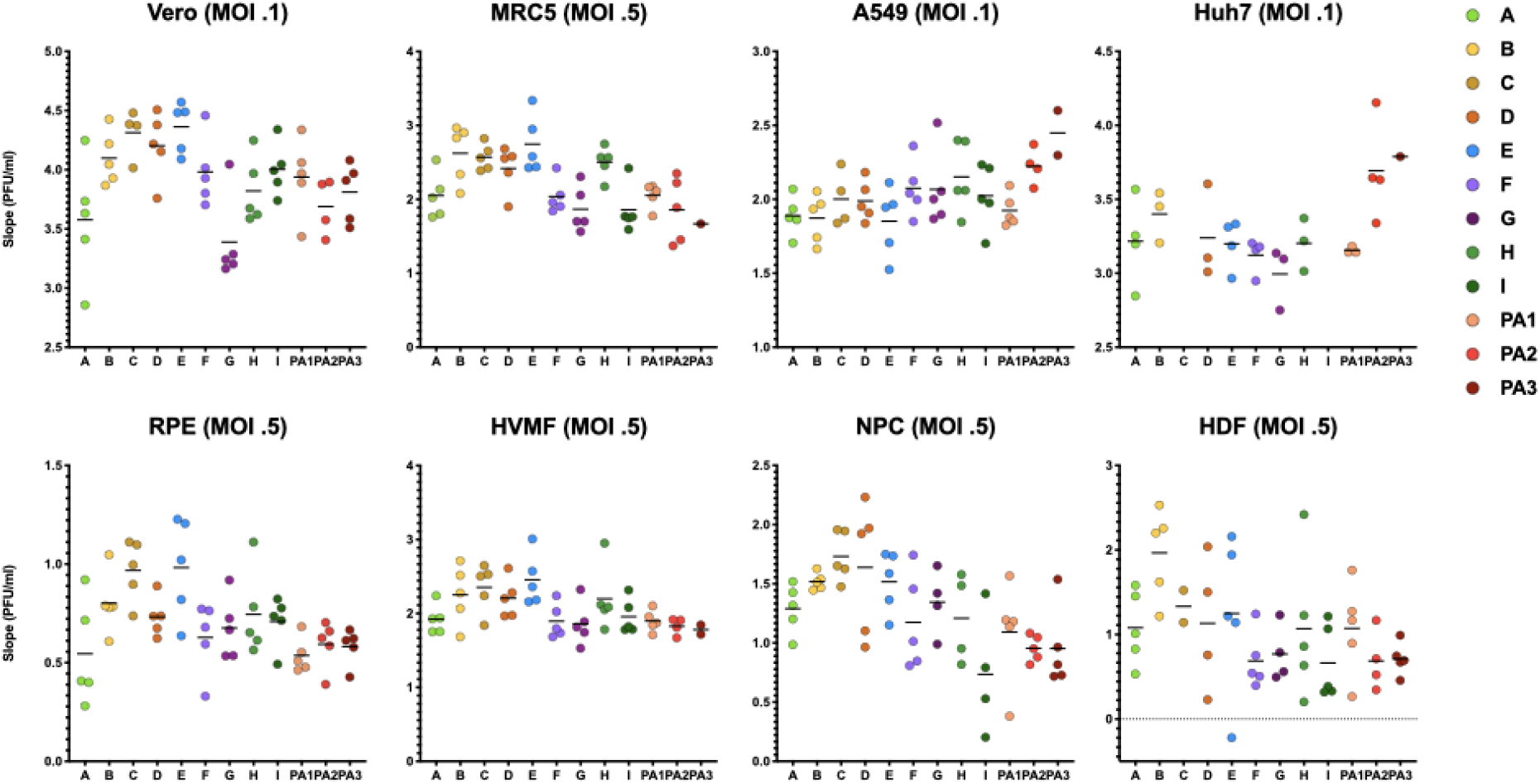
Zika virus replicative fitness growth rates. As a secondary endpoint to assess replicative fitness differences, we measured the growth rates of the replicative fitness curves across all 12 clades by fitting a line to every curve for all time points under exponential growth ^64^. Using this method, we are able to control for any variation in the infectious dose of the initial inocula. While there were no statistical differences found using this method, clades B, C, D, and E had the highest growth rates among primary human cells. Cell types are explained in (**Extended Data Table 2**).

**Extended Data Fig. 3.**
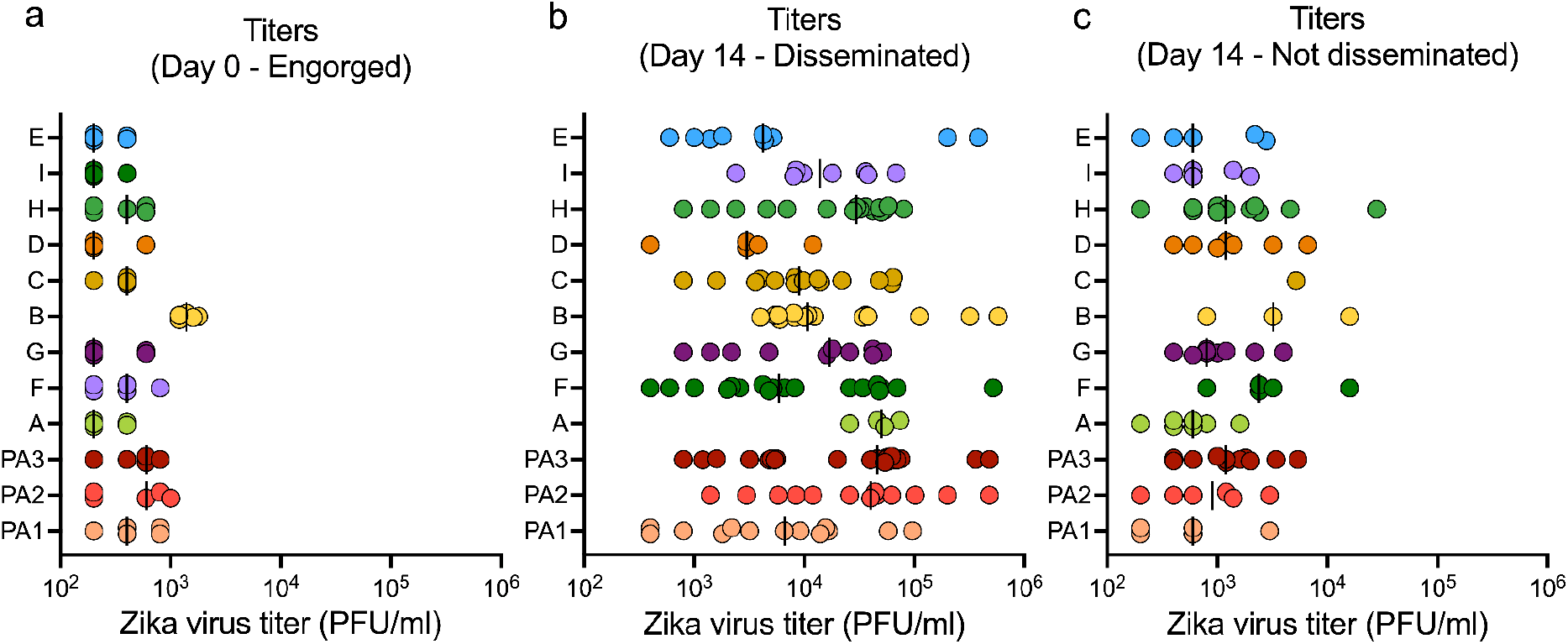
Zika virus titers in mosquito bodies after exposure via an infectious blood meal. We determined titers of the 12 Zika virus clades in *Aedes aegypti* (Poza Rica, Mexico) mosquito bodies. We determined Zika titers for (**a**) engorged mosquitoes at day 0, as well as (**b**) fully disseminated and (**c**) infected (but not disseminated) mosquitoes at day 14. Dots represent titers for individual mosquito bodies, and bars indicate the median. Letters indicate significance tested with a Kruskal-Wallis test, followed by a post-hoc Dunn’s test adjusted with false discovery rate (FDR) correction for multiple comparisons.

**Extended Data Fig. 4.**
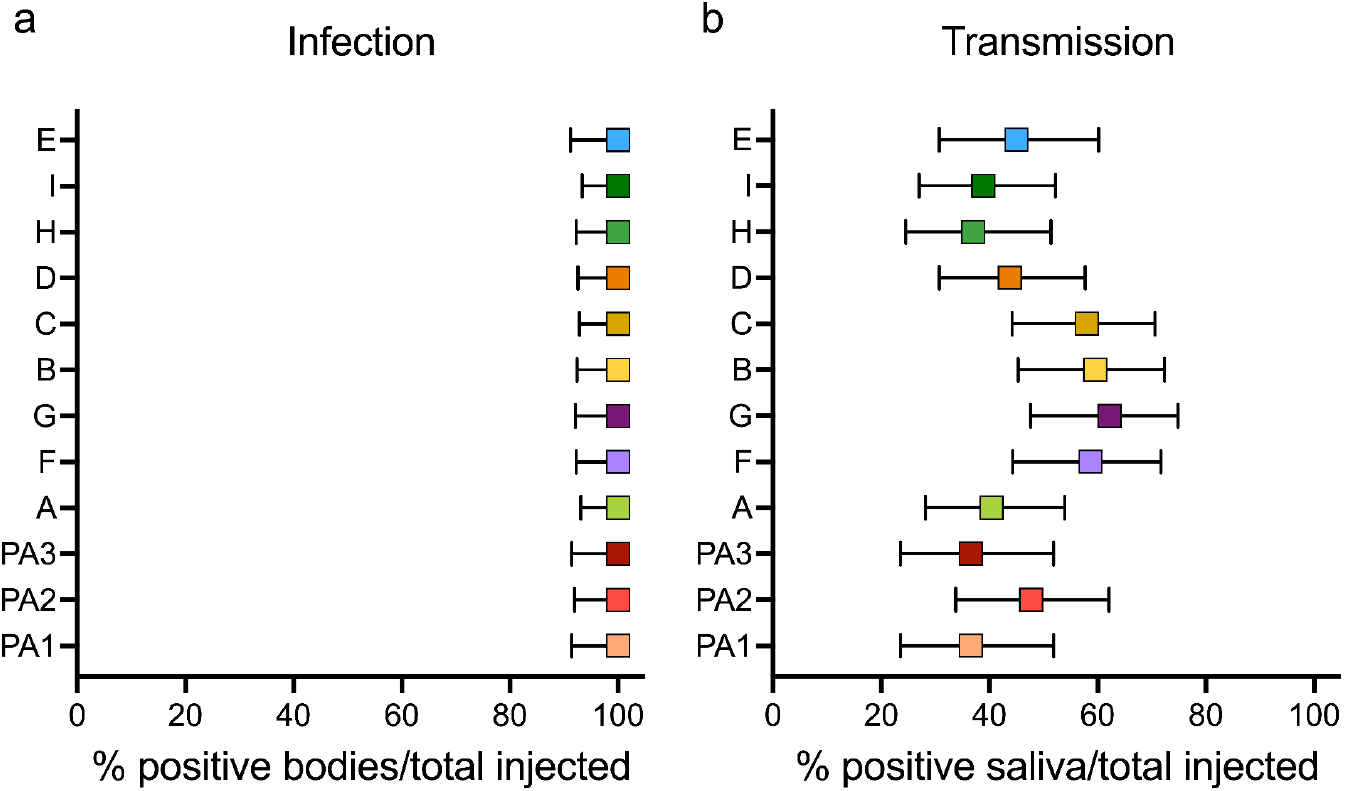
Evaluation of Zika virus fitness in *Aedes aegypti* mosquitoes infected by intrathoracic injection. We injected *Aedes aegypti* (Poza Rica) females with one of the twelve clades. After seven days incubation, (**a**) all females were confirmed to be infected (Zika virus-positive body), and (**b**) percentage of females with Zika virus-positive saliva out of the total number of infected females per clade (transmission) were similar among clades. Squares represent the rates expressed as percentages and error bars depict the 95% confidence intervals. We used a Fisher’s exact test for independence to analyze the data, but no significant differences were found.

## References

1. Metsky, H. C. et al. Zika virus evolution and spread in the Americas. Nature 546, 411–415 (2017).

2. Grubaugh, N. D., Faria, N. R., Andersen, K. G. & Pybus, O. G. Genomic Insights into Zika Virus Emergence and Spread. Cell 172, 1160–1162 (2018).

3. Avelino-Silva, V. I. et al. Potential effect of Zika virus infection on human male fertility? Rev. Inst. Med. Trop. Sao Paulo 60, e64 (2018).

4. Parra, B. et al. Guillain-Barré Syndrome Associated with Zika Virus Infection in Colombia. N. Engl. J. Med. 375, 1513–1523 (2016).

5. Manangeeswaran, M. et al. ZIKA virus infection causes persistent chorioretinal lesions. Emerg. Microbes Infect. 7, 96 (2018).

6. Hoen, B. et al. Pregnancy Outcomes after ZIKV Infection in French Territories in the Americas. N. Engl. J. Med. 378, 985–994 (2018).

7. Rossi, S. L., Ebel, G. D., Shan, C., Shi, P.-Y. & Vasilakis, N. Did Zika Virus Mutate to Cause Severe Outbreaks? Trends Microbiol. doi:10.1016/j.tim.2018.05.007.

8. Weaver, S. C. Emergence of Epidemic Zika Virus Transmission and Congenital Zika Syndrome: Are Recently Evolved Traits to Blame? MBio 8, (2017).

9. Pierson, T. C. & Diamond, M. S. The emergence of Zika virus and its new clinical syndromes. Nature 560, 573–581 (2018).

10. Liu, Y. et al. Evolutionary enhancement of Zika virus infectivity in Aedes aegypti mosquitoes. Nature 545, 482–486 (2017).

11. Shan, C. et al. A Zika virus envelope mutation preceding the 2015 epidemic enhances virulence and fitness for transmission. Proc. Natl. Acad. Sci. U. S. A. 117, 20190–20197 (2020).

12. Yuan, L. et al. A single mutation in the prM protein of Zika virus contributes to fetal microcephaly. Science 358, 933–936 (2017).

13. Jaeger, A. S. et al. Zika viruses of African and Asian lineages cause fetal harm in a mouse model of vertical transmission. PLoS Negl. Trop. Dis. 13, e0007343 (2019).

14. Tsetsarkin, K. A., Vanlandingham, D. L., McGee, C. E. & Higgs, S. A single mutation in chikungunya virus affects vector specificity and epidemic potential. PLoS Pathog. 3, e201 (2007).

15. Diehl, W. E. et al. Ebola Virus Glycoprotein with Increased Infectivity Dominated the 2013-2016 Epidemic. Cell 167, 1088–1098.e6 (2016).

16. Urbanowicz, R. A. et al. Human Adaptation of Ebola Virus during the West African Outbreak. Cell 167, 1079–1087.e5 (2016).

17. Volz, E. et al. Evaluating the Effects of SARS-CoV-2 Spike Mutation D614G on Transmissibility and Pathogenicity. Cell 184, 64–75.e11 (2021).

18. Washington, N. L. et al. Emergence and rapid transmission of SARS-CoV-2 B.1.1.7 in the United States. Cell 184, 2587–2594.e7 (2021).

19. Planas, D. et al. Reduced sensitivity of SARS-CoV-2 variant Delta to antibody neutralization. Nature (2021) doi:10.1038/s41586-021-03777-9.

20. Korber, B. et al. Tracking Changes in SARS-CoV-2 Spike: Evidence that D614G Increases Infectivity of the COVID-19 Virus. Cell 182, 812–827.e19 (2020).

21. Ohta, T. Synonymous and nonsynonymous substitutions in mammalian genes and the nearly neutral theory. J. Mol. Evol. 40, 56–63 (1995).

22. Benson, D. A. et al. GenBank. Nucleic Acids Res. 41, D36–42 (2013).

23. Edgar, R. C. MUSCLE: multiple sequence alignment with high accuracy and high throughput. Nucleic Acids Res. 32, 1792–1797 (2004).

24. Sagulenko, P., Puller, V. & Neher, R. A. TreeTime: Maximum-likelihood phylodynamic analysis. Virus Evol 4, vex042 (2018).

25. Nguyen, L.-T., Schmidt, H. A., von Haeseler, A. & Minh, B. Q. IQ-TREE: a fast and effective stochastic algorithm for estimating maximum-likelihood phylogenies. Mol. Biol. Evol. 32, 268–274 (2015).

26. Shannon, C. E. The mathematical theory of communication. 1963. MD Comput. 14, 306–317 (1997).

27. Tsetsarkin, K. A. et al. A Full-Length Infectious cDNA Clone of Zika Virus from the 2015 Epidemic in Brazil as a Genetic Platform for Studies of Virus-Host Interactions and Vaccine Development. MBio 7, (2016).

28. Dickinson, D. J., Ward, J. D., Reiner, D. J. & Goldstein, B. Engineering the Caenorhabditis elegans genome using Cas9-triggered homologous recombination. Nat. Methods 10, 1028–1034 (2013).

29. Hamel, R. et al. Biology of Zika Virus Infection in Human Skin Cells. J. Virol. 89, 8880–8896 (2015).

30. Li, C. et al. Zika Virus Disrupts Neural Progenitor Development and Leads to Microcephaly in Mice. Cell Stem Cell 19, 120–126 (2016).

31. El Costa, H. et al. ZIKA virus reveals broad tissue and cell tropism during the first trimester of pregnancy. Sci. Rep. 6, 35296 (2016).

32. Jurado, K. A. et al. Zika virus productively infects primary human placenta-specific macrophages. JCI Insight 1, (2016).

33. Luplertlop, N. et al. The impact of Zika virus infection on human neuroblastoma (SH-SY5Y) cell line. J. Vector Borne Dis. 54, 207–214 (2017).

34. Vicenti, I. et al. Comparative analysis of different cell systems for Zika virus (ZIKV) propagation and evaluation of anti-ZIKV compounds in vitro. Virus Res. 244, 64–70 (2018).

35. Gutiérrez-Bugallo, G. et al. Vector-borne transmission and evolution of Zika virus. Nat Ecol Evol 3, 561–569 (2019).

36. Weger-Lucarelli, J. et al. Vector competence of American mosquitoes for three strains of Zika virus. PLoS Negl. Trop. Dis. 10, e0005101 (2016).

37. Chouin-Carneiro, T. et al. Differential susceptibilities of *Aedes aegypti* and *Aedes albopictus* from the Americas to Zika virus. PLoS Negl. Trop. Dis. 10, e0004543 (2016).

38. Guo, X.-X. et al. *Culex pipiens quinquefasciatus*: a potential vector to transmit Zika virus. Emerg. Microbes Infect. 5, e102 (2016).

39. Main, B. J. et al. Vector competence of Aedes aegypti, Culex tarsalis, and Culex quinquefasciatus from California for Zika virus. PLoS Negl. Trop. Dis. 12, e0006524 (2018).

40. Fernandes, R. S. et al. Culex quinquefasciatus from Rio de Janeiro Is Not Competent to Transmit the Local Zika Virus. PLoS Negl. Trop. Dis. 10, e0004993 (2016).

41. Garcia-Luna, S. M. et al. Variation in competence for ZIKV transmission by Aedes aegypti and Aedes albopictus in Mexico. PLoS Negl. Trop. Dis. 12, e0006599 (2018).

42. Kryazhimskiy, S. & Plotkin, J. B. The population genetics of dN/dS. PLoS Genet. 4, e1000304 (2008).

43. Lemey, P., Minin, V. N., Bielejec, F., Kosakovsky Pond, S. L. & Suchard, M. A. A counting renaissance: combining stochastic mapping and empirical Bayes to quickly detect amino acid sites under positive selection. Bioinformatics 28, 3248–3256 (2012).

44. Harvey, W. T. et al. SARS-CoV-2 variants, spike mutations and immune escape. Nat. Rev. Microbiol. 19, 409–424 (2021).

45. Michlmayr, D., Andrade, P., Gonzalez, K., Balmaseda, A. & Harris, E. CD14+CD16+ monocytes are the main target of Zika virus infection in peripheral blood mononuclear cells in a paediatric study in Nicaragua. Nat Microbiol 2, 1462–1470 (2017).

46. Foo, S.-S. et al. Asian Zika virus strains target CD14+ blood monocytes and induce M2-skewed immunosuppression during pregnancy. Nat Microbiol 2, 1558–1570 (2017).

47. Kuo, L. et al. Reversion to ancestral Zika virus NS1 residues increases competence of Aedes albopictus. PLoS Pathog. 16, e1008951 (2020).

48. Lauring, A. S., Acevedo, A., Cooper, S. B. & Andino, R. Codon usage determines the mutational robustness, evolutionary capacity, and virulence of an RNA virus. Cell Host Microbe 12, 623–632 (2012).

49. Siedner, M. J., Ryan, E. T. & Bogoch, I. I. Gone or forgotten? The rise and fall of Zika virus. Lancet Public Health 3, e109–e110 (2018).

50. Hadfield, J. et al. Nextstrain: real-time tracking of pathogen evolution. Bioinformatics 34, 4121–4123 (2018).

51. Quick, J. et al. Multiplex PCR method for MinION and Illumina sequencing of Zika and other virus genomes directly from clinical samples. Nat. Protoc. 12, 1261–1276 (2017).

52. Grubaugh, N. D. et al. An amplicon-based sequencing framework for accurately measuring intrahost virus diversity using PrimalSeq and iVar. Genome Biol. 20, 8 (2019).

53. Pickett, B. E. et al. ViPR: an open bioinformatics database and analysis resource for virology research. Nucleic Acids Res. 40, D593–8 (2012).

54. O’Brien, J. D., Minin, V. N. & Suchard, M. A. Learning to count: robust estimates for labeled distances between molecular sequences. Mol. Biol. Evol. 26, 801–814 (2009).

55. Hasegawa, M., Kishino, H. & Yano, T. Dating of the human-ape splitting by a molecular clock of mitochondrial DNA. J. Mol. Evol. 22, 160–174 (1985).

56. Drummond, A. J., Ho, S. Y. W., Phillips, M. J. & Rambaut, A. Relaxed phylogenetics and dating with confidence. PLoS Biol. 4, e88 (2006).

57. Grubaugh, N. D. et al. Genomic epidemiology reveals multiple introductions of Zika virus into the United States. Nature 546, 401–405 (2017).

58. Mann, H. B. & Whitney, D. R. On a Test of Whether one of Two Random Variables is Stochastically Larger than the Other. Ann. Math. Stat. 18, 50–60 (1947).

59. Haynes, W. Benjamini–Hochberg Method. in Encyclopedia of Systems Biology (eds. Dubitzky, W., Wolkenhauer, O., Cho, K.-H. & Yokota, H.) 78–78 (Springer New York, 2013).

60. Virtanen, P. et al. SciPy 1.0: fundamental algorithms for scientific computing in Python. Nat. Methods 17, 261–272 (2020).

61. Seabold, S. & Perktold, J. Statsmodels: Econometric and statistical modeling with python. in Proceedings of the 9th Python in Science Conference (SciPy, 2010). doi:10.25080/majora-92bf1922-011.

62. Kruskal, W. H. & Wallis, W. A. Use of Ranks in One-Criterion Variance Analysis. J. Am. Stat. Assoc. 47, 583–621 (1952).

63. Waskom, M. seaborn: statistical data visualization. J. Open Source Softw. 6, 3021 (2021).

64. Wang, G. P. & Bushman, F. D. A statistical method for comparing viral growth curves. J. Virol. Methods 135, 118–123 (2006).

